# Assembly and glycosylation of *Helicobacter pylori* sheathed flagella

**DOI:** 10.64898/2025.12.30.697013

**Authors:** Rajeev Kumar, Shoichi Tachiyama, Huaxin Yu, Samira Heydari, Jiaqi Guo, Jack M. Botting, Wangbiao Guo, Timothy R. Hoover, Jun Liu

**Affiliations:** Department of Microbial Pathogenesis, Yale School of Medicine, New Haven, CT USA; Microbial Sciences Institute, Yale University, West Haven, CT USA; Department of Microbiology, University of Georgia, Athens, GA USA

**Keywords:** macromolecular assemblies, membranous sheath, motility, flagellin, glycosylation

## Abstract

The bacterial flagellum is a complex nanomachine essential for motility, colonization, and invasion in diverse species. *Helicobacter pylori* has evolved elaborate sheathed flagella that enable migration through the highly viscous gastric mucus layer to reach its colonization niche on the gastric epithelium, yet the molecular basis for these unique adaptations has remained elusive. Here, we use *in-situ* single particle cryo-electron microscopy to determine near-atomic structures of the flagellar filament within the membranous sheath of *H. pylori*. The major flagellin FlaA constitutes the bulk of the filament, whereas the minor flagellin FlaB contributes critically to the hook-proximal region. Both FlaA and FlaB form a conserved core surrounded by variable surface-exposed domains. Our structures further reveal that pseudaminic-acid glycans decorate these domains, where they mediate inter- and intra-subunit contacts that stabilize the filament and confer a negatively charged surface. Together, these findings support a model in which the filament rotates independently of the membranous sheath to drive *H. pylori* motility and provide a molecular framework for understanding how the sheathed flagellum enables colonization and persistence within the gastric niche.

**Significance Statement:** We present the first *in-situ* near-atomic structure of the sheathed flagellar filament in *Helicobacter pylori*, revealing distinctive adaptations that underpin the pathogen’s unique motility and persistent infection. Our *in-situ* structures show that the two flagellins, FlaA and FlaB, assemble into an extended and exceptionally stable filament through an extensive hydrogen-bonding network. Pseudaminic acid glycans decorate the surface-exposed domains, where they stabilize inter-subunit packing and render the surface negatively charged and hydrophilic. These findings, which provide insight into the assembly of the flagellar filament and its relationship to the surrounding sheath, provide a structural framework for developing strategies to disrupt *H. pylori* motility and infection.

## Introduction

*Helicobacter pylori* is a gastric bacterium that colonizes approximately half the world’s population (1, 2). Having coexisted with humans for at least 50,000 years (3), *H. pylori* has evolved remarkable strategies to evade the immune system and colonize its preferred niche, the gastric mucosa. Although *H. pylori* colonization does not cause symptoms in most people, it can trigger chronic stomach inflammation and significantly increase the risk of developing peptic ulcer disease, chronic gastritis, and gastric cancer (4–6). The ability of *H. pylori* to traverse the gastric mucous layer and colonize the underlying epithelial cell surface is promoted by its characteristic helical shape (7, 8), urease production (9, 10), and robust motility in viscous media (10–12). *H. pylori* motility is driven by a bundle of unipolar flagella, each of which is enclosed by a membranous sheath that is continuous with the outer membrane. Although the role of the sheath has yet to be determined experimentally, possible functions include protecting the flagella from dissociating in the gastric acidic environment, facilitating adherence, and avoiding surveillance by the host innate immune system (13).

The bacterial flagellum has been extensively characterized in model organisms such as *Escherichia coli* and *Salmonella enterica*. It is composed of three principal parts: the rotary motor, the hook, and the filament (14). The motor is embedded in the cell envelope and is responsible for flagellar assembly and rotation. At least 20 different proteins are involved in building the core components of the motor, including the MS-ring, C-ring, LP-rings, rod, export apparatus, and torque-generating stator units. The hook functions as a universal joint, transmitting torque from the rod to the filament (15, 16). The rotating filament serves as a propeller to drive bacterial motility.

Although the core components of bacterial flagella are broadly conserved, the sheathed flagella of *H. pylori* have evolved distinctive adaptations that enable motility within and colonization of the gastric mucosa. The *H. pylori* genome encodes numerous motor accessory proteins that are structurally and functionally distinct from those of enteric bacteria and are essential for motility (17–20). Furthermore, the *H. pylori* flagellar filament differs from that of enteric species in that it is assembled from two flagellins, FlaA and FlaB, which do not to activate Toll-like receptor 5, contributing to immune evasion and persistent colonization (21–23). FlaA constitutes the majority of the filament, whereas the minor flagellin FlaB localizes at the filament base near the hook (24). Consistent with their distinct roles, deletion of *flaA* results in truncated filaments and severely impaired motility, whereas *flaB* mutants produce full-length filaments with moderately reduced motility (25). Intriguingly, both FlaA and FlaB undergo extensive O-glycosylation with 5,7-diacetamido-3,5,7,9-tetradeoxy-L-glycero-L-manno-nonulosonic acid (Pse5Ac7Ac), a nine-carbon pseudaminic acid derivative structurally related to the sialic acid Neu5Ac found on human cell surfaces. Disruption of flagellin glycosylation abolishes filament assembly and motility in *H. pylori* (26). Although the precise functional role of glycosylation remains incompletely understood, it has been proposed to stabilize inter-subunit interactions during filament assembly (27).

Here, we deploy *in-situ* single particle cryo-EM to determine near-atomic structures of the sheathed flagellar filament in *H. pylori*, revealing how two flagellins, FlaA and FlaB, assemble the filament through highly conserved domains that form in the central core of the filament and variable domains located on the surface of the filament. We further identify residues within the variable domains of FlaA and FlaB that are glycosylated, as well as protein-ligand interactions that likely promote filament assembly and stability.

## Results

### High-resolution *in-situ* structures of the sheathed flagellar filament

FlaA is the main component of the filament whereas FlaB is a minor flagellin found near the hook (24). To elucidate the unique adaptations of the sheathed flagellum in *H. pylori* (Fig. 1A), we employed an *in-situ* single-particle cryo-EM approach to determine the filament structures directly from frozen-hydrated bacteria, without filament isolation or purification (Fig. 1). Using cryoSPARC (28), filaments were first automatically selected and processed to generate 2D class averages (Fig. 1D; SI Appendix, Fig. S1). The majority of 2D classes display an intact bilayer membranous sheath surrounding the filament, with filaments lacking the sheath also observed (Fig. 1D; SI Appendix, Fig. S1A). The spacing between the filament and sheath varies from approximately 1 to 4 nm for majority of 2D classes (Fig. 1D and SI Appendix, Fig. S1A). Filaments enclosed by the sheath were selected for subsequent helical reconstruction and refinement, yielding a 2.65 Å resolution structure of the sheathed filament (Fig. 1E, F and SI Appendix, Fig. S1B-D, Fig. S2A-C). To determine the filament structures near the filament tip and the hook, we manually selected filament segments and determined their *in-situ* structures at 3.19Å and 3.22Å resolution, respectively (SI Appendix, Fig. S2). All three filament structures exhibit nearly identical helical parameters: a helical twist of 65.40° and rise of 4.68 Å (SI Appendix, Table S1).

**Figure 1.**
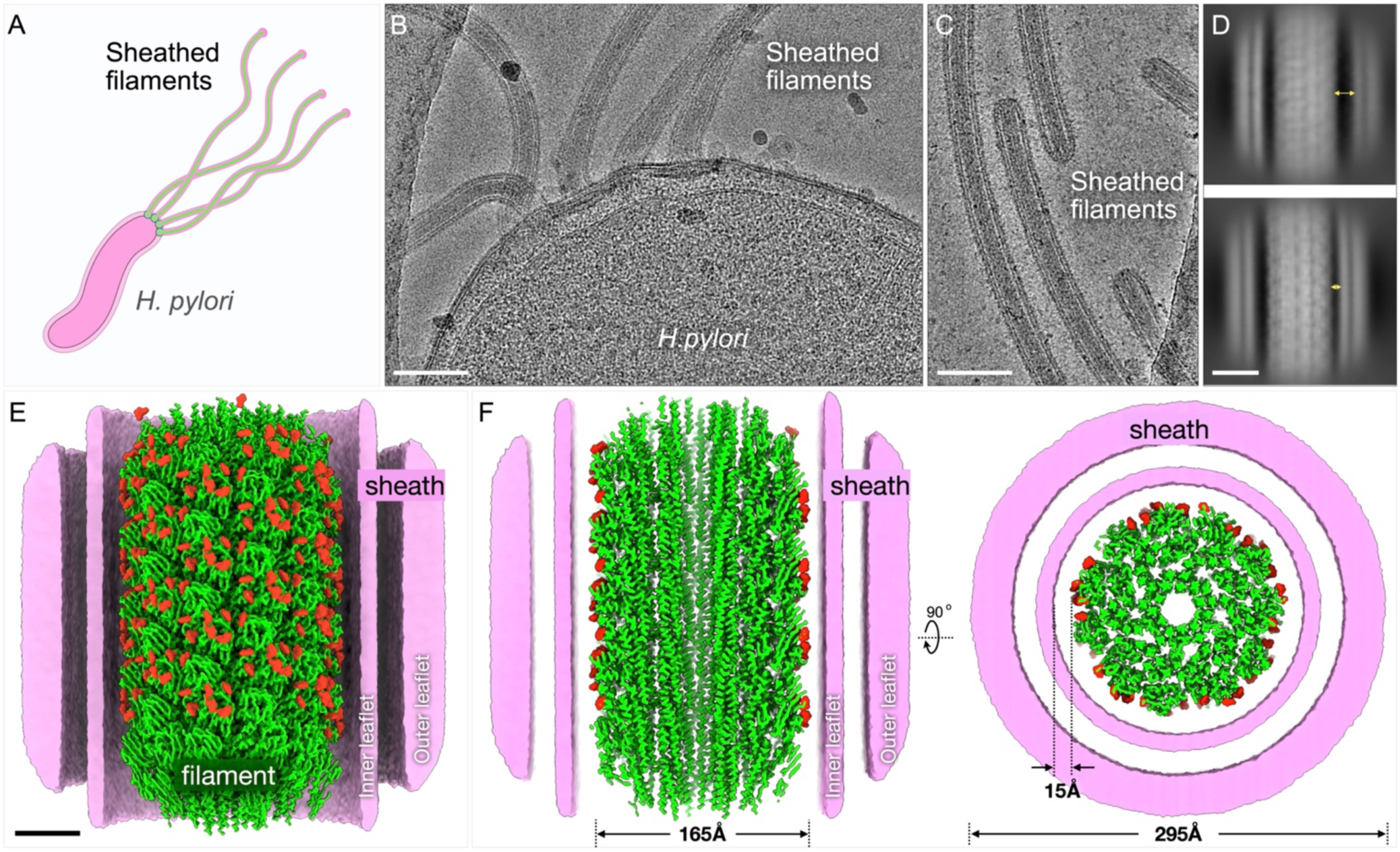
*In-situ* cryo-EM structure of the sheathed flagellar filament at near-atomic resolution. **(A)** A schematic model of a *H. pylori* cell with multiple sheathed flagella. (**B)** A typical cryo-EM micrograph collected from a cell pole containing multiple sheathed flagella. Scale bar is 100nm. (**C**) A typical cryo-EM micrograph collected from the filament tips. (**D**) 2D class averages of the flagellar filaments show variable distance (1-4nm) between the flagellar filament and the membranous sheath. Sale bar is 10nm. (**E**) A surface rendering of the flagellar filament structure at 2.65 Å resolution with a helical symmetry of twist 65.40° and rise 4.68 Å. A cut-through view of the membranous sheath enables a better view of the filament, the inner and outer leaflets of the membranous sheath. The filament is colored in green and glycans are colored in red. Sale bar is 4nm. (**F**) Left panel: A cross-sectional view of the filament shows the filament core. The diameter of the filament is 165 Å. The averaged diameter of the membranous sheath is 295 Å. (**F**) Right panel: A top view of the sheathed flagella shows 11 protofilaments and the surrounding membranous sheath.

Consistent with flagellar filaments of *S. enterica* and other bacteria (29, 30), the *H. pylori* flagellar filament forms an 11-protofilament tubular structure composed of helical arrays of flagellin subunits (Fig. 1E-G). The filament diameter is ∼165 Å and the central lumen measures ∼28 Å in diameter (Fig. 1F; SI Appendix, Fig. S1, S2). Including the surrounding membranous sheath, the total diameter of the sheathed flagellum reaches ∼295 Å (Fig. 1F). Importantly, our *in-situ* structures at near-atomic resolution afforded us the opportunity to analyze unique features of the sheathed flagellar filament in intact bacteria.

### The *in-situ* structure of the major flagellin FlaA

Two flagellins, FlaA and FlaB, share 58% amino acid identity and 75% similarity across their entire lengths (24) (SI Appendix, Fig. S3A). AlphaFold-predicted flagellin structures show a canonical feature where a single flagellin peptide folds to form a helical coiled-coil structure and beta sheets and can be classified into the typical four domains: D0, D1, D2, and D3 (SI Appendix, Fig. S3B)(31, 32). Domains D0 and D1 are highly conserved whereas the surface-exposed domains are more divergent (SI Appendix, Fig. S3C). To test which flagellin is present in our maps near the tip and middle regions of the filament we fit the predicted FlaA and FlaB models into our cryo-EM density maps, followed by manual model building in Coot (33) and refinement in Phenix (34). Our data showed that the refined FlaA model fit well into the densities corresponding to the tip and central regions of the filament, while the FlaB model did not (SI Appendix, Fig. S4A, B). Specifically, residues Arg-23, Ser-26, Arg-66, Thr-237, and Arg-294 in FlaB did not fit the density appropriately, whereas the corresponding FlaA residues – Asn-23, Lys-26, Ala-66, Arg-227, and Glu-284 – aligned well with the map (SI Appendix, Fig. S4A, B).

Our cryo-EM structures confirm that FlaA consists of four major domains: D0, D1, D2, and D3 (Fig. 2A-C). The D0 domain is composed of ND0 (residues 1–36) and CD0 (residues 472–510), and D1 includes ND1 (residues 37–167) and CD1 (residues 412–472) (Fig. 2C). Assembly of FlaA subunits within the filament revealed that Arg-381 and Glu-219 form bifurcated hydrogen bonds, while Glu-146 and Arg-418 engage in inter-chain hydrogen bonding with neighboring protomers (Fig. 2D). These interactions form an extensive hydrogen-bond network that is also found in other bacterial filaments and is critical for filament assembly and structural stability (Fig. 2D) (30, 35).

**Figure 2.**
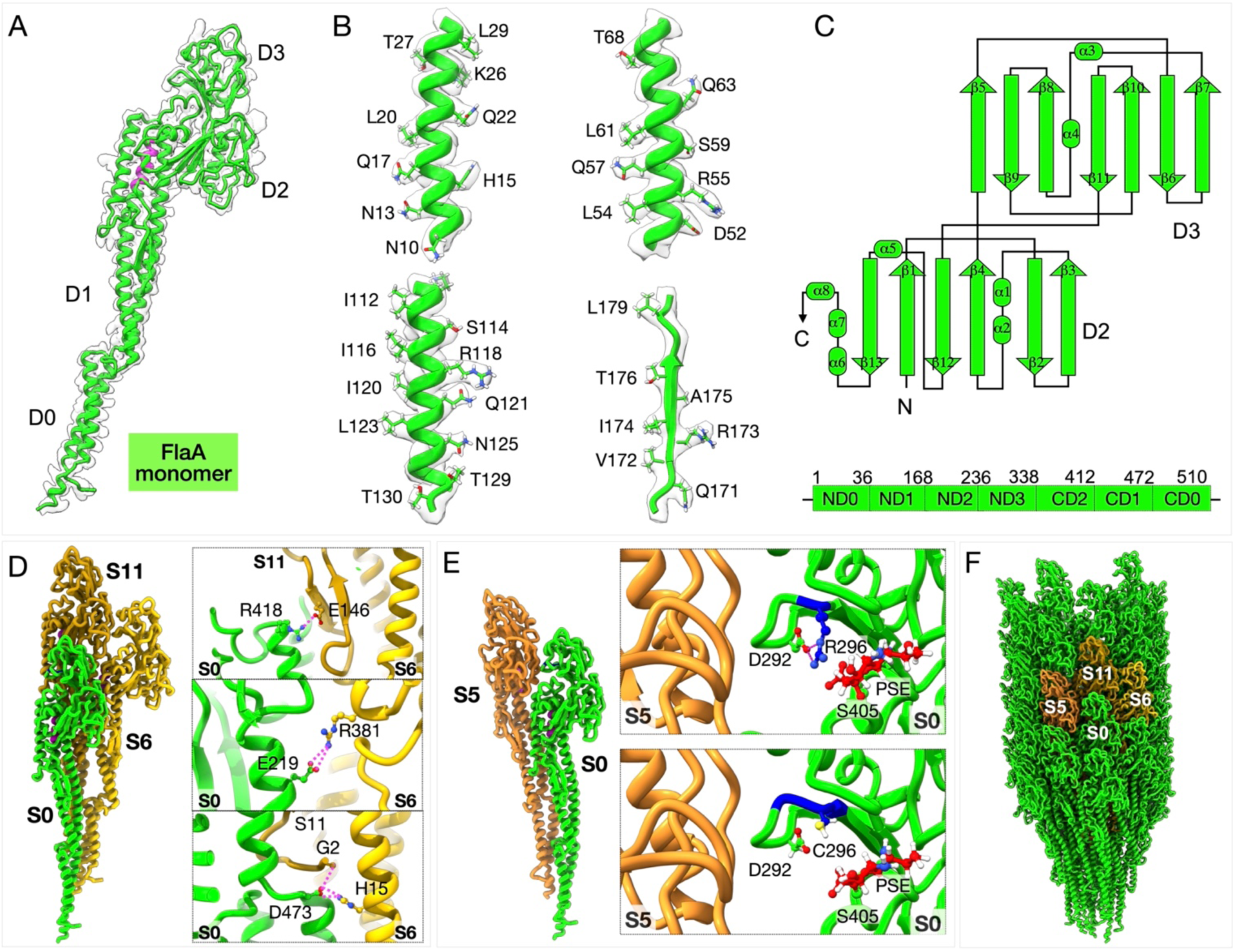
Structure of major flagellin FlaA from *H. pylori*. (**A**) Refined model of the FlaA monomer fitted into the cryo-EM density map. (**B**) Representative density fittings for the D0 and D2 domains of FlaA. (**C**) Schematic representation of the D2 and D3 domains (top) and linear arrangement of the four FlaA domains (bottom). Domains D2 and D3 adopt immunoglobulin-like folds. (**D**) Interactions between adjacent FlaA monomers (S0, S6, and S11). The right panels highlight key residues involved in hydrogen-bond interactions; left, side view of the filament model; right, top view. (**E**) Point mutation in domain D3 of *H. pylori* B128 FlaA (Cys-296→Arg-296, blue). The top panel shows Arg-296 forming a hydrogen bond with Asp-292, whereas no such interaction occurs in the wild-type Cys-296 (bottom). (**F**) Side view of the FlaA filament. Three adjacent FlaA monomers (S5, S6, and S11) are colored in orange.

The *in-situ* structure of FlaA further reveals that residue 296 is an arginine (Fig. 2E; SI Appendix, Fig. S5A), which is different from a cysteine in the original sequence of *H. pylori* B128 FlaA (36). DNA sequencing of *flaA* from the *H. pylori* B128 strain used in this study confirmed that codon 296 encodes arginine (CGC) rather than cysteine (TGC) (SI Appendix, Fig. S5A). Comparative analysis of FlaA sequences from different *H. pylori* strains – including 26695, J99, SS1, G27, and ATCC 43504 – showed that all possess arginine at position 296 (SI Appendix, Fig. S5B). Notably, the *H. pylori* 7.13 strain, a mouse-adapted derivative of B128, also contains Arg-296 and exhibits a significantly larger swim halo in soft-agar assays than its parental strain (36). Although it remains unclear whether the Cys-to-Arg substitution contributes to enhanced motility of *H. pylori* 7.13 compared to its parental strain, our *in-situ* structure indicates that Arg-296 forms a hydrogen bond with Asp-292 (Fig. 2E), potentially contributing to filament stability and function.

### Distinct flagellin surface glycosylation sites in *H. pylori*

Post-translational modifications of bacterial flagellins, particularly glycosylation, are critical for motility in many bacterial species (35, 37). Mass spectrometry analysis of FlaA from *H. pylori* strain 1061 previously identified glycosylation by pseudaminic acid (Pse5Ac7Ac) at seven serine and threonine residues within the D2 and D3 domains (26). Consistent with these findings, fitting the refined FlaA model into the *in situ* cryo-EM density map derived from middle regions of the filament revealed additional densities at seven sites near specific serine and threonine residues on the surface of the variable D2 and D3 domains (Fig. 3A, B). Five of these glycosylated residues – Thr-181, Ser-246, Ser-354, Thr-364, and Ser-405 – match those identified by mass spectrometry (SI Appendix, Fig. S6A, B) (26). The remaining two glycosylated residues, Ser-207 and Ser-393, are adjacent to previously reported sites (Ser-208 and Ser-395) (Fig. 3B; SI Appendix, Fig. S6B). This discrepancy unlikely results from sequence heterogeneity, as the amino acid sequences surrounding Ser-207 and Ser-393 are identical between FlaA from *H. pylori* strains B128 and 1061 (used in the mass spectrometry analysis) (SI Appendix, Fig. S6B).

**Figure 3.**
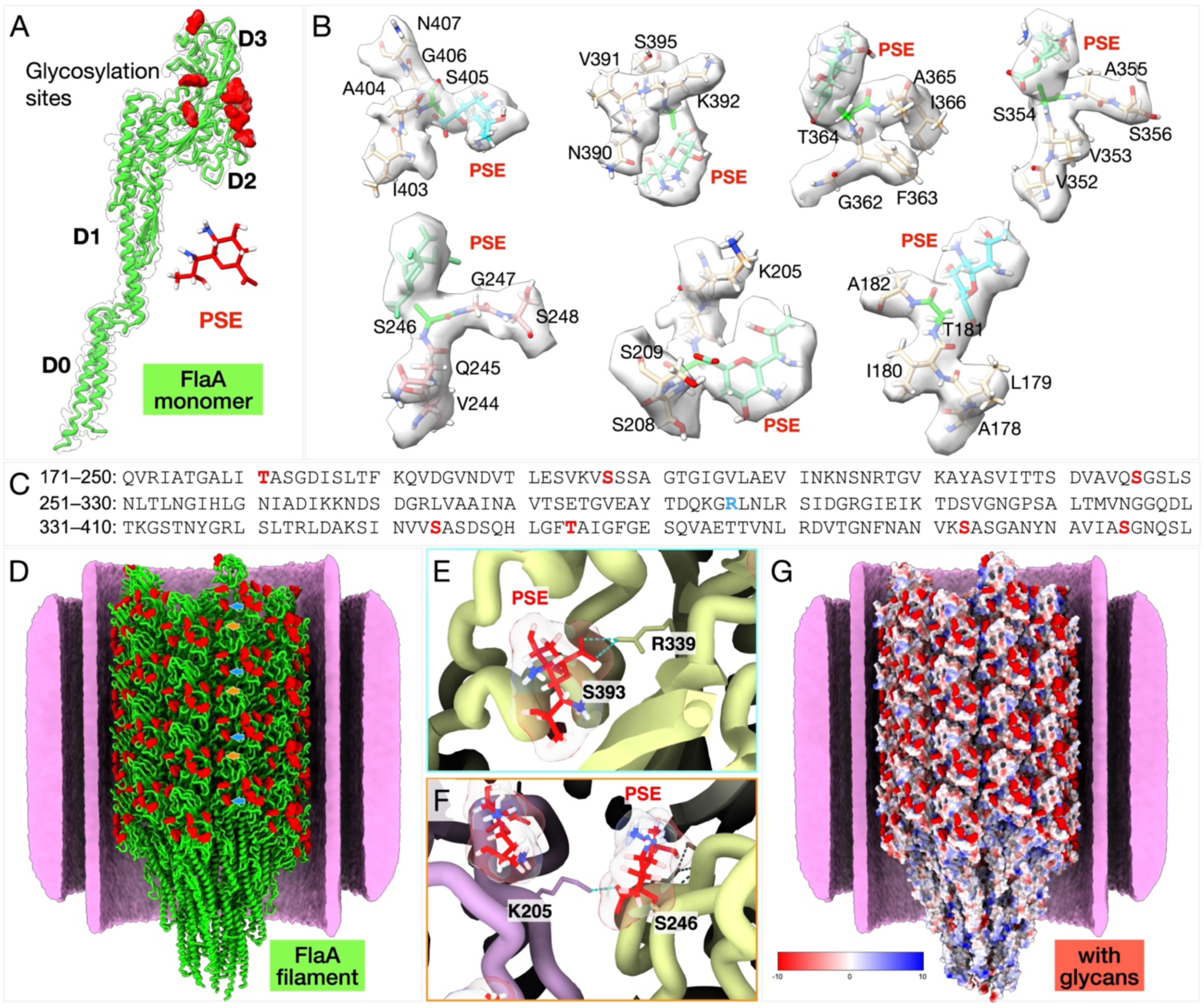
The FlaA filament surface is extensively decorated with glycans. (**A**) The FlaA monomer contains seven glycosylation sites located on domains D2 and D3. (**B**) Model of Pse5Ac7Ac fitted into glycan densities, showing O-linked glycosylation via serine and threonine residues. (**C**) Sequence of the FlaA D2 and D3 domains, with glycosylation sites colored in red and Arg-296 in blue. (**D**) Ribbon model of the FlaA filament showing extensive glycan densities on the filament surface (red blobs). Representative image showing formation of H-bond by Pse5Ac7Ac within the same protofilament **(E)** and with the neighboring protofilament **(F)**. (**G**) Electrostatic surface representations reveal an overall negative surface charge of the *H. pylori* flagellar filament, primarily due to glycosylation.

The *in-situ* structure further revealed that several Pse5Ac7Ac glycans form hydrogen bonds with neighboring amino acid side chains, suggesting a stabilizing role in filament architecture (SI Appendix, Fig. S6A). For example, the glycan attached to Ser-393 in D2 forms hydrogen bonds with Asn-407, Asn-398, and Arg-339, while the glycan at Ser-246 in D3 forms a hydrogen bond with Lys-205 in D2 of an adjacent protofilament (SI Appendix, Fig. S6C, D). These interactions likely contribute to filament stability and inter-subunit packing. Importantly, the glycan decorations render an overall negative surface charge to the filament (Fig. 3F). Two of the major phospholipids in *H. pylori*, cardiolipin and phosphatidylglycerol (38), are anionic phospholipids, and the negatively charge surface of the filament imparted by the glycan decorations may electrostatically repel these negatively charged phospholipids within the inner leaflet in the membranous sheath.

### FlaB structure and distribution in *H. pylori*

We further determined a cryo-EM structure derived from the filaments near the hook region and built a FlaB model (Fig. 4A-F; SI Appendix, Fig. S2G-I). The overall domain organization of FlaB resembles that of FlaA, comprising four domains (D0-D3) (Fig. 4B). Although the D0 and D1 domains of FlaA and FlaB share high structural similarity, distinct differences were observed within the D2 and D3 domains, which constitute the variable, surface-exposed regions (Fig. 4E).

**Figure 4.**
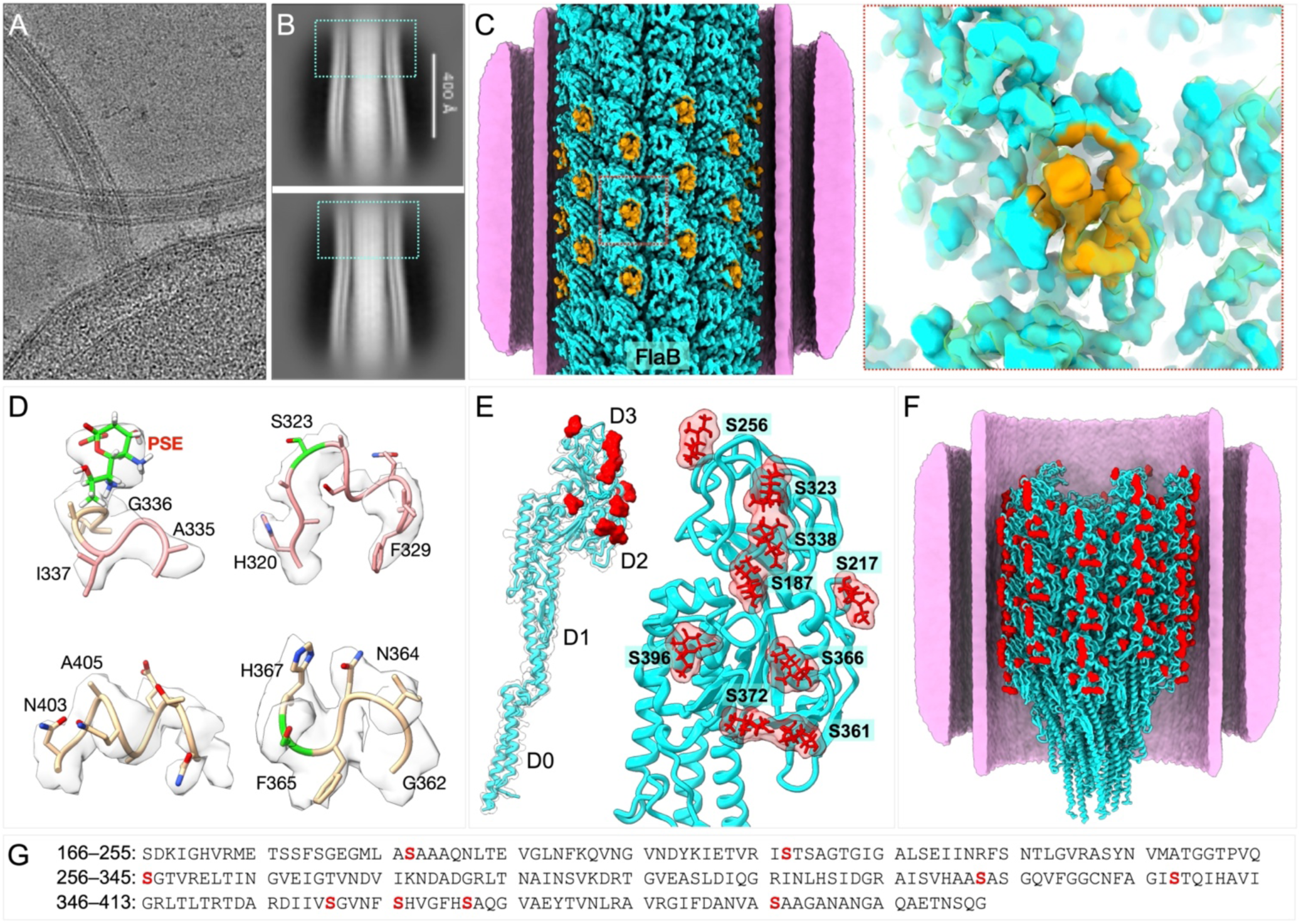
Structure of FlaB and its distinct glycosylation sites. (**A**) Cryo-EM image from the cell tip showing the FlaB filament near the hook region. (**B**) Representative 2D class averages from near-hook regions. The blue boxes are the FlaB filament regions. (**C**) Surface rendering of the FlaB filament structure. The orange region shows a distinct loop in FlaB. (**D**) Representative density fittings of the D2 and D3 domains of FlaB with glycans. (**E**) Cryo-EM structure reveals nine glycosylation sites located on domains D2 and D3 of FlaB. (**F**) Ribbon model of the FlaB filament showing extensive glycan densities on the filament surface (red blobs). (**G**) Sequence of FlaB D2–D3 domains, with glycans colored in red.

Notably, the D2 and D3 domains of FlaB contain clear densities corresponding to nine O-linked glycosylation sites modified by Pse5Ac7Ac (Fig. 4E; SI Appendix, Fig. S7). Earlier work identified ten Pse5Ac7Ac-modified sites in FlaB by mass spectrometry (26) though the precise positions could not be determined due to the predominance of FlaA in those samples. The glycosylated tryptic peptides and number of attached Pse5Ac7Ac units, however, were quantified. Except for one discrepancy, the number of glycosylation sites resolved in our *in-situ* FlaB structure closely match those inferred from the mass spectrometry data. The difference involves the peptide spanning Met-173 to Lys-200, which was previously reported to contain two Pse5Ac7Ac units (26) but in our cryo-EM map only a single glycosylation site (Ser-187) was observed (Fig. 4E).

Four of the FlaB glycosylation sites lie within regions of greater sequence homology to FlaA than to the rest of the D2 and D3 domains (SI Appendix, Fig. S7A-C). While FlaA and FlaB share 43% identity and 66% similarity across the entire D2/D3 domains, the regions surrounding their common glycosylation sites exhibit 58% identity and 84% similarity (SI Appendix, Fig. S7A). Structural alignment of the D2 and D3 domains of FlaA and FlaB yielded an RMSD of 0.513 Å, indicating strong structural conservation despite sequence divergence (SI Appendix, Fig. S7D).

Overall, our near-atomic *in-situ* cryo-EM structure of FlaB defines its domain architecture and pinpoints glycosylation sites, highlighting the power of *in-situ* single-particle cryo-EM in resolving post-translational modifications in complex assemblies.

## Discussion

Using *in-situ* single-particle cryo-EM, we determined near-atomic structures of the sheathed flagellar filament, revealing molecular adaptations that underlie the unique motility and persistent infection of *H. pylori*. FlaA forms the bulk of the filament, while FlaB localizes proximally near the hook, consistent with previous observations (24). Both flagellins share conserved D0/D1 domains and variable D2/D3 domains (Fig. 2, 4). The D0/D1 domains form the filament core and the central channel for unfolded subunit secretion (39), whereas the D2/D3 domains comprise the sheath-proximal outer surface (Fig. 3A; Fig. 4A).

Our *in-situ* structures identify these glycosylation sites (Fig. 3-5) and reveal Pse5Ac7Ac-mediated hydrogen bonding that stabilizes the filament. In FlaA, Ser-207 and Ser-393 are glycosylated, differing slightly from the predicted Ser-208 and Ser-395; this discrepancy likely reflects peptide overlap during mass-spectrometric analysis. In FlaB, nine glycosylation sites were observed, compared with ten previously predicted (26). This difference may stem from strain-specific variation in deglycosylation by Cds6 (HP0518), an L,D-carboxypeptidase implicated in peptidoglycan trimming and cell-shape control (40, 41). Flagellins from a *cds6* mutant are hyper-glycosylated, containing roughly three-fold more Pse5Ac7Ac than those from wild type (37). These results suggest that FlaA and FlaB are glycosylated in the cytoplasm before export, after which Cds6 removes excess Pse5Ac7Ac units. Strain-dependent Cds6 activity may therefore explain variations reported for the number of Pse5Ac7Ac units associated with FlaB in the previous mass-spectrometry analysis (25) and the *in-situ* structure of FlaB.

**Figure 5.**
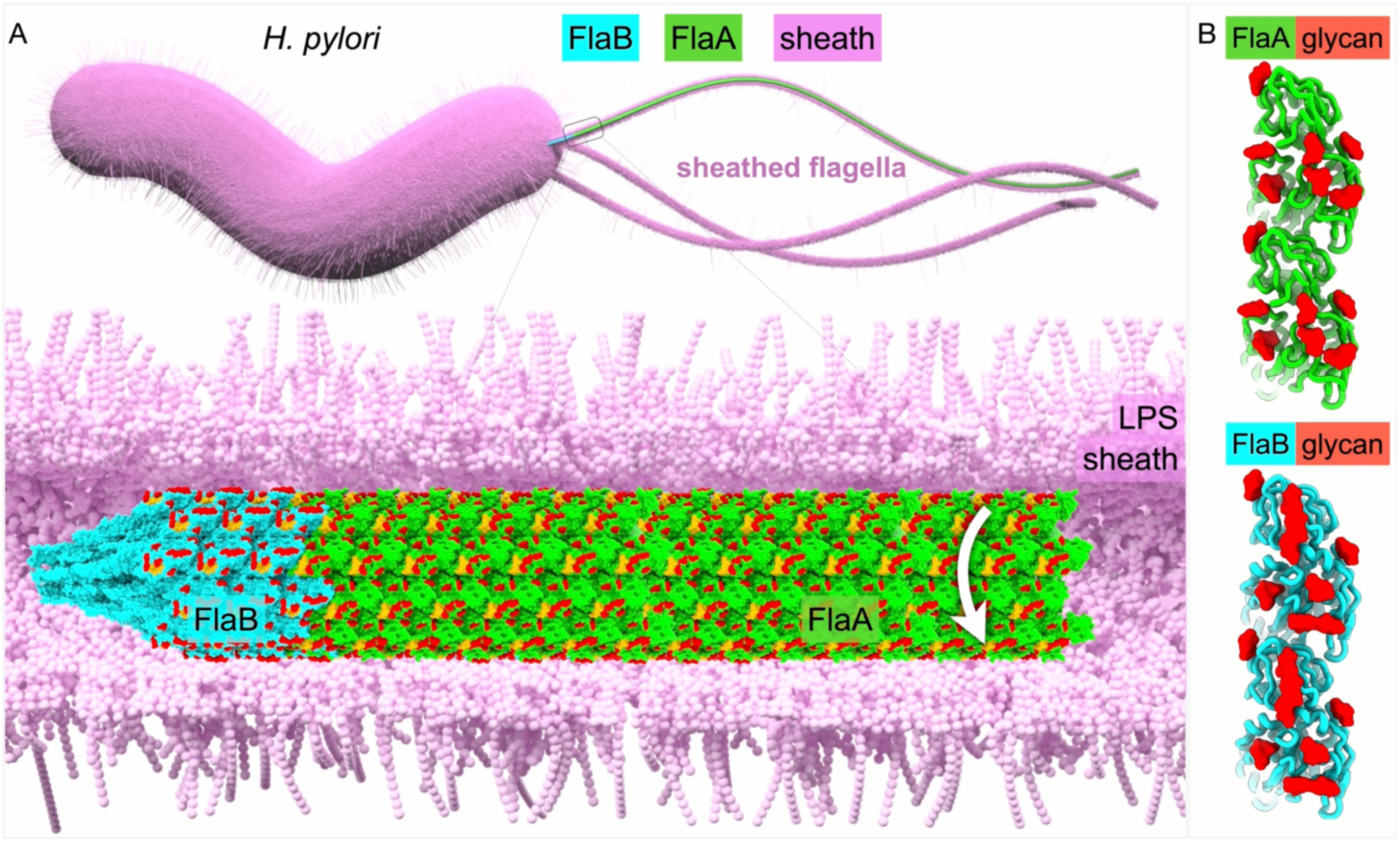
Model of assembly and rotation of the sheathed flagellar filament. (**A**) Top: Model of an *H. pylori* cell possessing multiple sheathed flagella, each comprising a motor, hook, and filament. The filament consists primarily of the major flagellin FlaA, with the minor flagellin FlaB located at the hook-proximal region. Bottom: Atomic models of the FlaB and FlaA filaments, both extensively covered by glycans. The while arrow indicates that the filament could rotate independently within the surrounding membranous sheath. (**B**) Comparison of distinct surface glycans decorating FlaA and FlaB.

We hypothesize that flagellin glycosylation directly contributes to filament-sheath dynamics. Decoration of FlaA and FlaB with Pse5Ac7Ac introduces a dense array of negative charges on the filament surface, which likely repels the negatively charged phospholipid headgroups of the inner leaflet of the membranous sheath (Fig. 5). This electrostatic repulsion supports a model in which the flagellar filament rotates independently from the surrounding membranous sheath (42). Consistent with this model, 2D class averages reveal that the sheath is not tightly bound to the filament but separated by a variable gap (Fig. 1D; SI Appendix, Fig. S1A). Comparable electrostatic interactions were recently observed in *V. cholerae* sheathed flagellum (43), although the filament–sheath spacing is substantially narrower than in *H. pylori* (SI Appendix, Fig. S8). Interestingly, unlike *H. pylori*, *V. cholerae* flagellins are not glycosylated, but the filament surface is enriched in acidic residues (PDB 9N8B), suggesting that many bacterial species have evolved distinct strategies to generate a negatively charged, hydrophilic flagellar filament surface for optimal rotation against the surrounding membranous sheath.

An intriguing question is why does *H. pylori* employ two flagellin species to assemble the flagellar filament? *H. pylori* FlaB is dispensable for the formation of functional flagellar filaments, although the *H. pylori flaB* mutant displays reduced motility compared to wild type (25). It is possible that the higher number of glycosylation sites in FlaB than FlaA increases the stability of the filament, allowing *H. pylori* to better navigate viscous environments. A recent study that generated a cryo-EM structure of the *S. enterica* FlgK-FlgL hook-filament junction proposed that the hook-filament junction protects the filament from mechanical stress emanating from the flexible hook (44). The portion of the *H. pylori* flagellar filament that is formed by FlaB may be more resistant to any mechanical stress not buffered by the hook-filament junction and thereby help stabilize the filament. Further analysis of *H. pylori* FlaB and its relationships with FlaA and hook-filament junction proteins will be needed to understand the specific roles of FlaB in the assembly and function of the flagellar filament, as well as the role of glycosylation in FlaB function.

In summary, our *in-situ* near-atomic structures of the *H. pylori* flagellar filament reveal a highly specialized motility apparatus. Similar to many flagellar filaments from different bacterial species, the *H. pylori* filament is stabilized by a conserved, canonical hydrogen-bonding network and inter-domain contacts. Unique to *H. pylori*, extensive Pse5Ac7Ac glycosylation reinforces filament integrity, modulates electrostatic properties, and supports a model of an independent filament rotation within a flexible sheath. Together, these unique properties of the *H. pylori* flagellar filament likely equip the bacterium to endure the harsh gastric environment and sustain persistent colonization.

## Materials and Methods

### Bacterial culture preparation

*H. pylori* B128 was grown on tryptic soy agar (TSA) plate with 30 μg/mL of kanamycin and 5% heat-inactivated horse serum at 37°C in 10% CO_2_ atmosphere for two days. Then, the strain was inoculated and grown in Brucella broth with the antibiotics and 10% heat inactivated fetal bovine serum at 37°C with shaking in a reduced O_2_ condition using CampyGen 2.5 L for the overnight. Next day, the overnight bacterial culture was inoculated into a fresh medium and grown for four hours. The bacterial culture was centrifuged at 1,500 ×g for 10 min, and bacterial pellets were resuspended into phosphate-buffered saline (PBS) at pH 7.4 and adjusted OD_600_ of 0.8.

### Cryo-EM sample preparation

To prepare sample for the single particle cryo-EM experiment, 5μL of the bacterial sample was deposited on fresh glow-discharged cryo-EM grids (Quantifoil, Cu, 200 mesh). Grids were blotted for 6–8 seconds under 90% humidity and rapidly plunge-frozen in liquid propane – ethane mixture ethane using a GP2 plunger (Leica).

### Single particle cryo-EM data collection

The grids of frozen-hydrated specimens were loaded into a Titan Krios electron microscope (Thermo Fisher Scientific) equipped with a field emission gun, K3 summit detector, and GIF BioQuantum imaging filter (Gatan). SerialEM (45) was used to collect data from bacterial flagellar filaments at magnification 81,000× with the physical pixel size of 1.068 Å and defocus ranging from −1.0 to −2.0μm. In total, 50,562 movies were collected in dose fraction mode with a dose of 73 e⁻/Å² distributed over 40 frames per movie.

### Cryo-EM image processing and 3D structure reconstruction

All data processing was performed using CryoSPARC (28). Beam-induced image drift was corrected using patch motion correction, and the contrast transfer function (CTF) was estimated with patch CTF estimation. Filament tracer was utilized to select 1,070,759 particles, initially extracted from 4× binned images (4.272 Å/pixel), followed by 2D classification to remove non-filament particles. Selected particles were then re-extracted at the original pixel size (1.068 Å/pixel) and used for ab initio reconstruction to create an initial model for iterative heterogeneous refinement, further eliminating low-quality particles. These particles underwent iterative non-uniform refinement without imposed symmetry, followed by local refinement, achieving a 3.26 Å resolution reconstruction. A symmetry search determined helical parameters with a rise of 4.675 Å and a twist of 65.45°. Using 228,410 high-quality particles, helical refinement produced a 3D reconstruction at 2.80 Å resolution without cyclic symmetry, and final non-uniform refinement yielded a map at 2.65 Å resolution (later identified as FlaA through modeling). From the initial 2D classification, a filament-like class with a varying diameter was noted and attributed to the filament–hook junction. The volume alignment tool shifted particle centers to the near-pole flagellar filament after determining a 3D structure through homogeneous refinement. The resulting 60,029 particles were re-extracted at the original pixel size (1.068 Å/pixel), and subsequent homogeneous and helical refinements resolved the structure at 3.22 Å resolution without cyclic symmetry (later identified as FlaB through modeling), sharing the same helical parameters as FlaA with a rise of 4.675 Å and a twist of 65.45°.

A total of 2,830 particles from the flagellar tip were manually picked and extracted from 2× binned images (2.136 Å/pixel) for *ab initio* reconstruction to generate initial models. Non-uniform refinement was performed, followed by symmetry expansion using parameters determined for FlaA and FlaB, yielding 30,715 particles. The FlaA density map was imported, low pass filtered to 10 Å and used as the reference volume for multiple rounds of local refinement. Particle centers were shifted along the flagellar tip at 48.06 Å intervals using Volume Alignment Tools. The resulting 275,229 particles were re-extracted at the original pixel size (1.068 Å/pixel) and subjected to additional rounds of local refinement, achieving a final resolution of 3.19 Å.

### Model building and refinement

Since *H. pylori* flagella consists of two flagellins FlaA and FlaB (24). We retrieved the FlaA and FlaB flagellin sequence of *H. pylori* B128 from JGI IMG database (46) (IMG ID 2877310586 and 2877311207 for FlaA and FlaB, respectively). We used these sequences to generate AlphaFold models (31, 32). The AlphaFold monomer structures for FlaA were initially docked into the Cryo-EM density map using UCSF ChimeraX (47). This docked model was manually fitted in Coot (33) for the mismatched residue and then Real space refined using Phenix (34). This step was repeated until the best fitted model was generated and the model quality was validated by Molprobity (48). A similar approach was used to build an atomic model for flagellin FlaB. Although the overall structures of FlaA and FlaB are similar, the difference lies on the surface exposed loop region (SI Appendix, Fig. S7D). This minor loop difference was prominent enough to distinguish between FlaA and FlaB. Our next step was to build the complete filament structure fragment for each density map. Hence, we docked the refined FlaA and FlaB monomer model against the cryo-EM density maps in a step wise manner (*i.e.,* one at a time) followed by real space refinement of flagellin structures in Phenix and model building in Coot. The final filament structure was achieved by Real space refinement in Phenix (34), and the model was validated in Molprobity (48). Similar protocol was followed for each map corresponding to tip area, middle area and the near hook area. The refinement statistics are provided in SI Appendix, Table S1. UCSF ChimeraX (47) was used to visualize the cryo-EM maps and the atomic models.

### Sequencing of B128 *flaA*

Genomic DNA was prepared from the B128 strain using the QIAamp DNA kit (Qiagen). PCR primer pair 5’-ATGGCTTTTCAGGTCAATACAAATATCAATGC-3’ (flaA-F) and 5’-CTAAGTTAAAAGCCTTAAGATATTTTGTTGAACGG-3’ (flaA-R) together with Q5 High fidelity DNA polymerase (NEB) were used to amplify the entire coding region of *flaA* from the *H. pylori* B128 gDNA. The resulting amplicon was submitted to the Keck DNA Sequencing Facility (Yale University) for Sanger sequencing using the flaA-F and flaA-R primers.

## Data availability

The refined atomic structure coordinates of reconstructed *H. pylori* flagellin FlaA and FlaB was deposited to the Protein Data Bank (PDB) with accession code 9YGU for FlaA and 9YH1 for FlaB and the cryo-EM map was deposited under EMDB cryo-EM database with accession code EMD-72941 and EMD-72948, respectively.

## Acknowledgments

We thank Jennifer Aronson for critical reading and editing of the manuscript. R.K., S.T., H.Y., S. H., W.G., and J.L. were partly supported by grants R01AI189907, R01AI087946 and R01AI132818 from National Institute of Allergy and Infectious Diseases (NIAID) and National Institutes of Health (NIH). Cryo-EM data were collected at the Yale Cryo-EM resources funded in part by the NIH grant 1S10OD023603-01A1. We thank the Keck DNA Sequencing Facility at Yale for their assistance with sequencing.

## Supplemental Information

**SI Appendix, Fig. S1.**
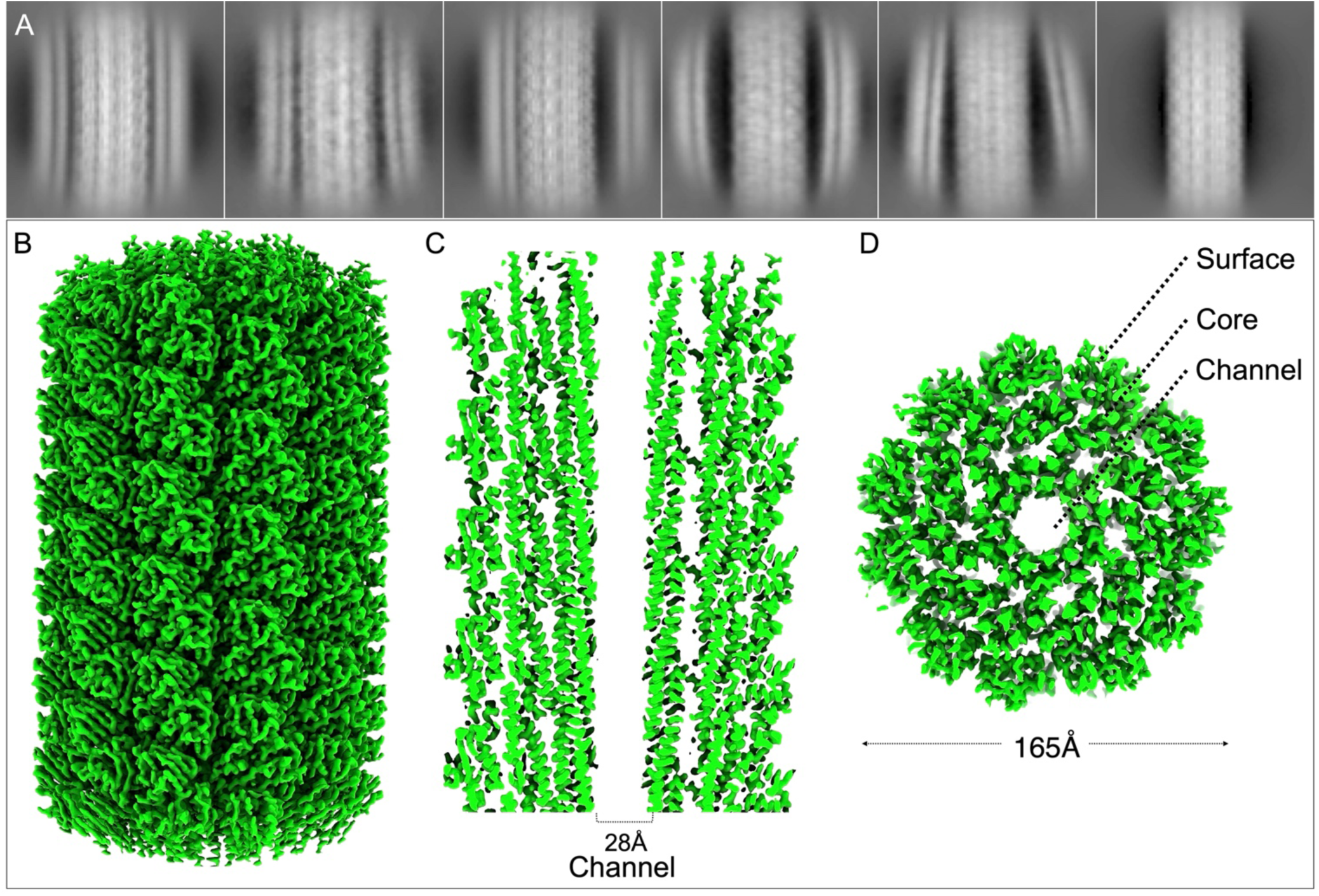
Cryo-EM reveals the flagellar filament structure enclosed by a membranous sheath with a variable diameter in *H. pylori*. **(A)** 2D class averages showing the presence of the membranous sheath around the filament. Notably, the spacing between the filament and sheath ranges from 1 to 4nm. No sheath is present in one class. (**B**) 3D surface view of the FlaA filament. (**C**) A vertical cross section of the filament shows filament core and central channel. (**D**) A horizontal cross section of the filament shows the central channel and surface-exposed domains.

**SI Appendix, Fig. S2.**
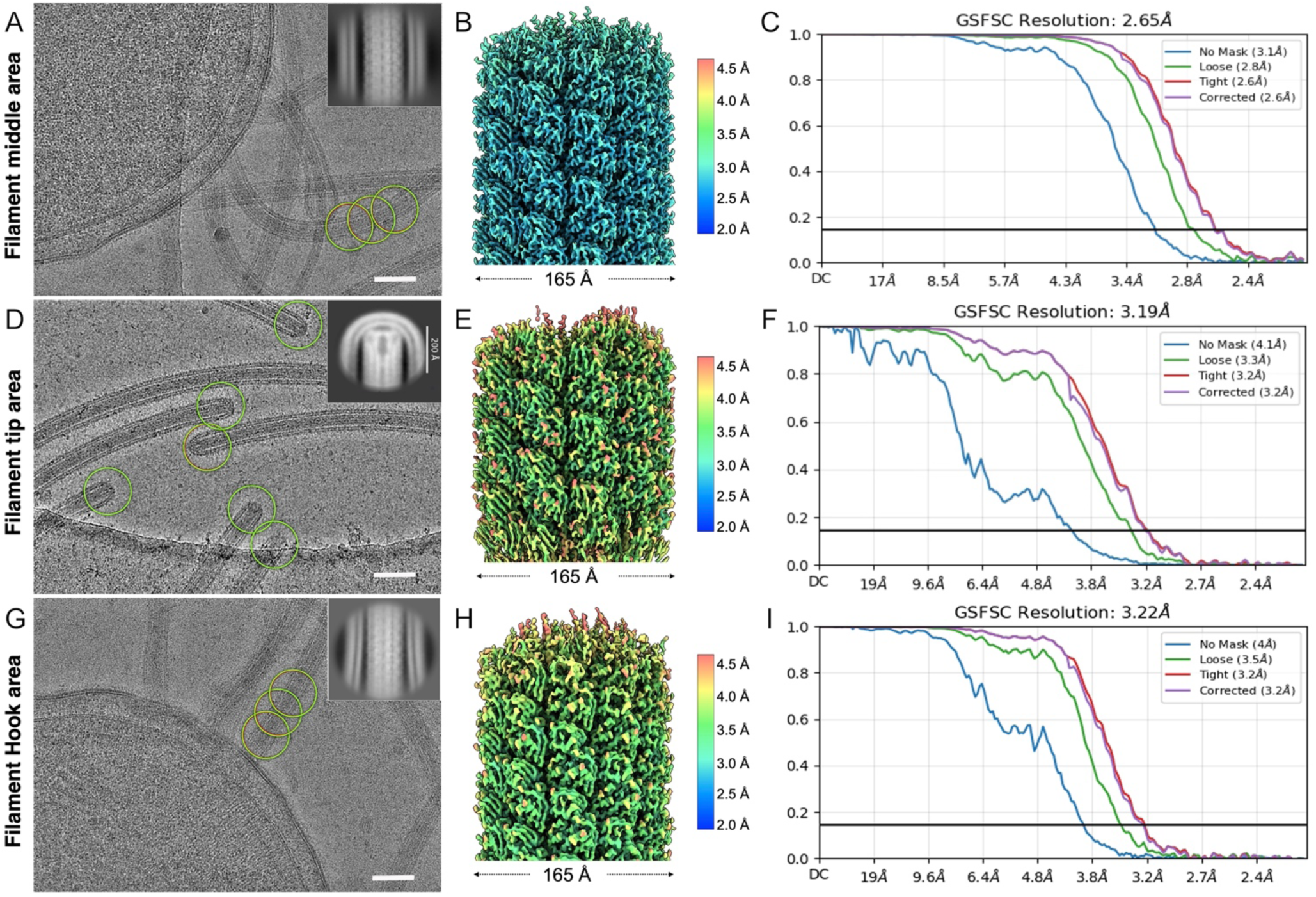
Cryo-EM data processing and resolution evaluation. (**A**, **D**, **G**) Micrographs and selected particles (green circles) of flagellar filaments at the middle (filament tracer auto-picking), tip and near-hook areas (manual picking) with representative 2D classes in inset. scale bars are 50 nm. (**B**, **E**, **H**) Local resolution maps from the middle, tip, and near-hook region, respectively. (**C**, **F**, **I**) Corresponding FSC resolution plots of the structures shown in panels **B**, **E**, and **H**, respectively.

**SI Appendix, Fig. S3.**
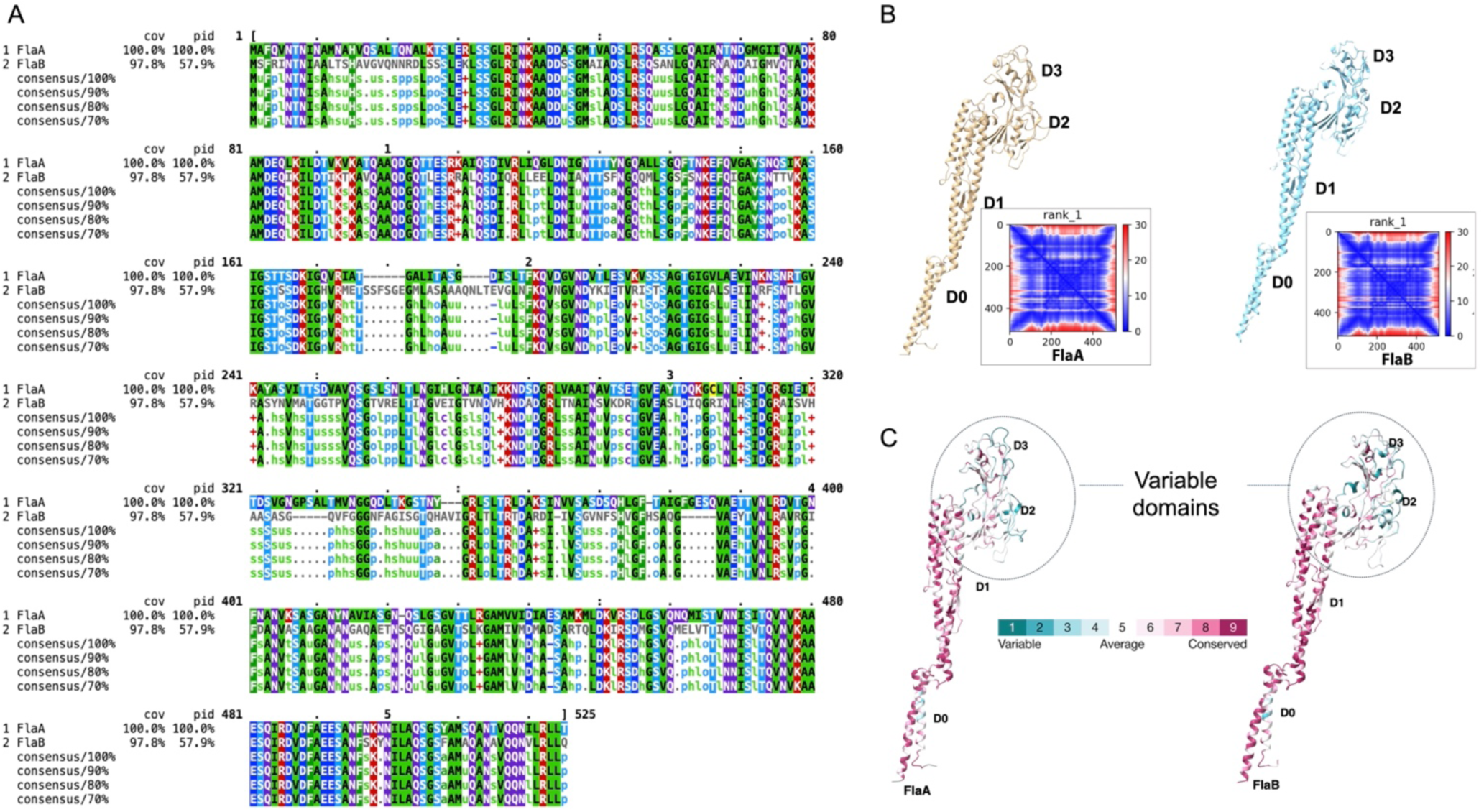
Flagellin FlaA and FlaB in *H. pylori* are conserved. (**A**) Sequence alignment of FlaA and FlaB is represented using Mview (49). (**B**) AlphaFold-predicted structures of flagellins FlaA and FlaB with very high confidence (31, 32). (**C**) Analysis of sequence conservative nature of flagellins FlaA and FlaB using Consurf web server (50).

**SI Appendix, Fig. S4.**
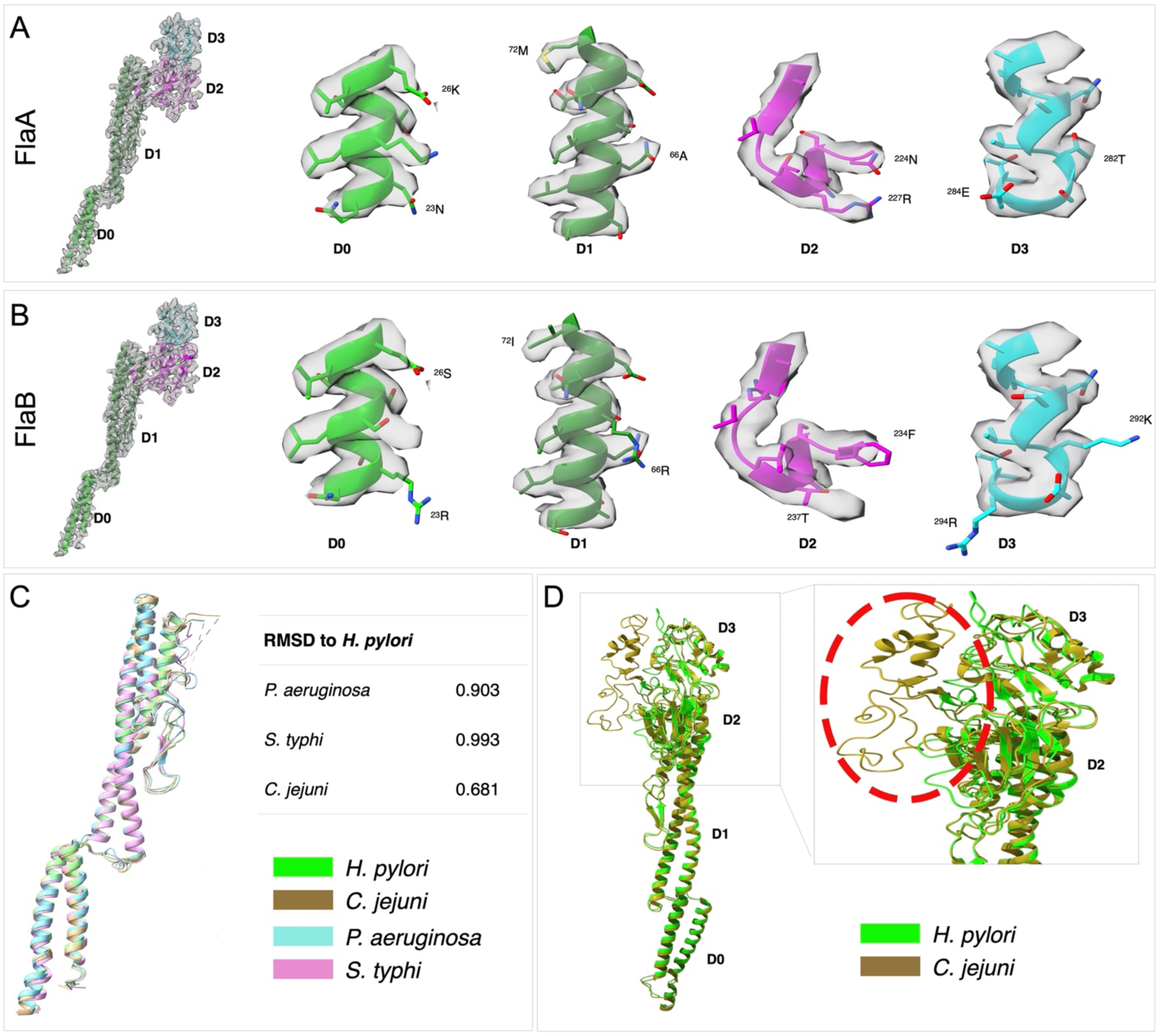
Comparison of FlaA structure from *H. pylori* with flagellin structures from other bacteria. (**A**) Left: Ribbon diagram of the refined structure of FlaA fitted into the cryo-EM density map. Right: Zoom-in views of the fitting at different domains of FlaA. (**B**) Left: Ribbon diagram of the refined structure of FlaB fitted into the cryo-EM density map. Right: Zoom-in views of the fitting at different domains of FlaB. Note that some side chains do not fit well into the densities compared to the fitting of FlaA. (**C**) Comparison of D0-D1 domains of *H. pylori* FlaA with those of *P. aeruginosa* FliC, *S. typhi* FliC, and *C. jejuni* FliA. (**D**) Comparison of FlaA structures from *H. pylori* and *C. jejuni*. *C. jejuni* has extra D4 domain which is absent in *H. pylori*.

**SI Appendix, Fig. S5.**
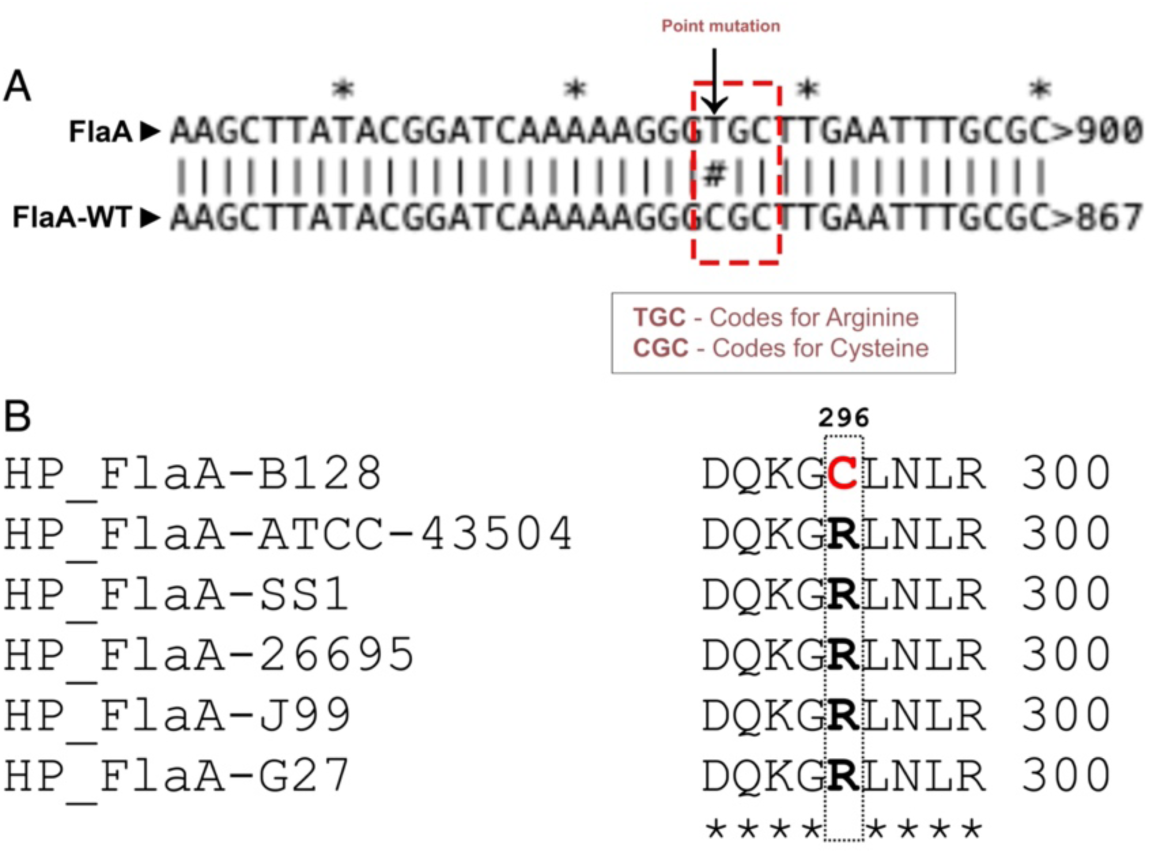
Sequencing of flagellin FlaA in *H. pylori* B128. **(A)** Sequencing confirms the presence of arginine at position 296 in FlaA representing a missense mutation in the *H. pylori* B128 *flaA* sequence in the NCBI and JGI IMG databases. (**B)** Image showing sequence alignment of FlaA from *H. pylori* strains-26695, J99, SS1, G27, and ATCC 43504, indicating presence of arginine residues at 296 position is common in these strains while the wild type *H. pylori* B128 has cysteine at same position. The star represents perfect match of residue.

**SI Appendix, Fig. S6.**
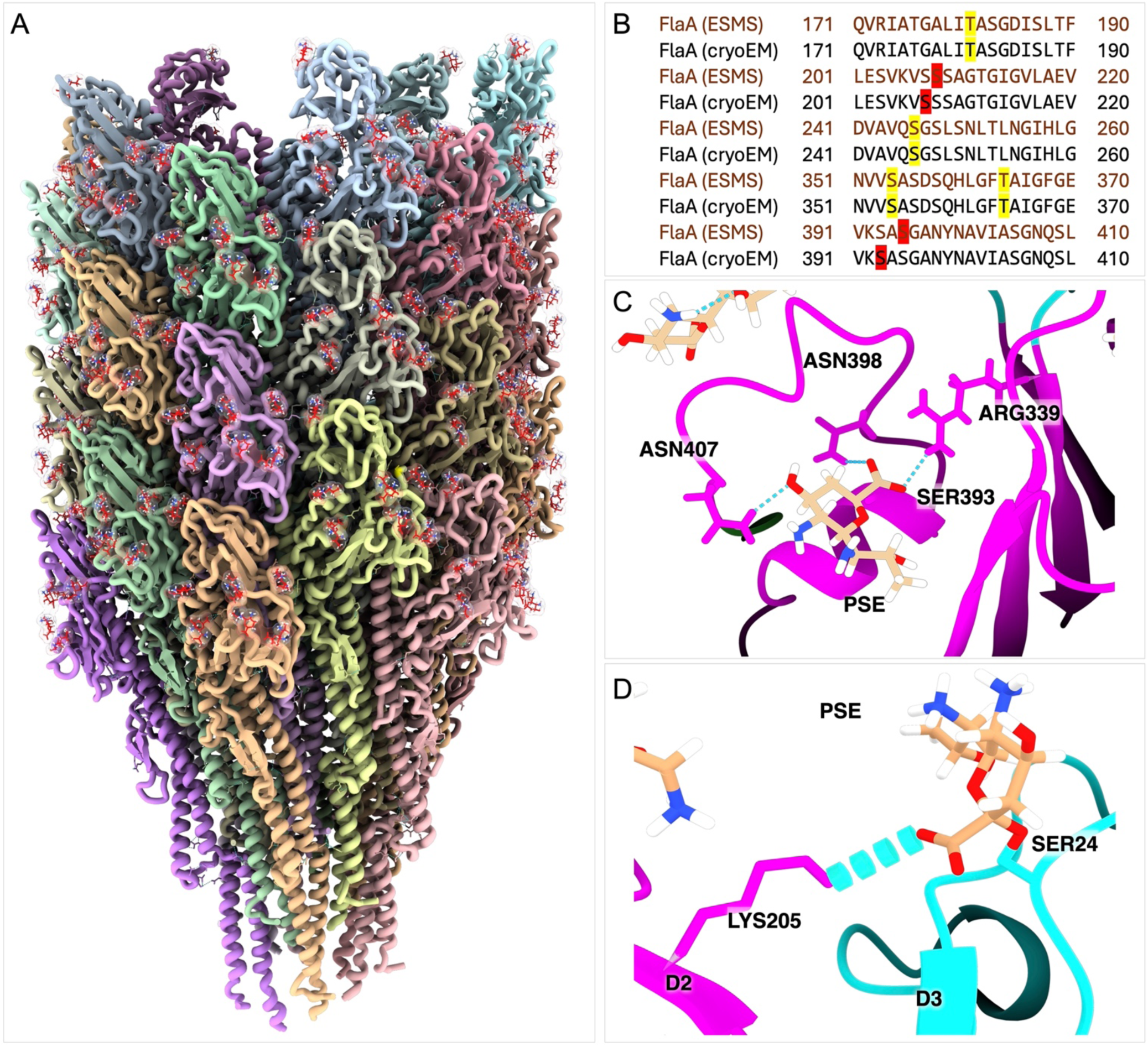
Glycosylation sites on flagellin FlaA in *H. pylori* B128. (**A**) The FlaA filament is decorated with extensive glycans. (**B**) Comparison of the glycosylation sites on flagellin FlaA revealed by cryo-EM and those previously identified by ESMS (26). (**C**) Zoomed image of Ser-393 residue participating in glycosylation forms an H-bond with adjacent charged residues in FlaA. (**D**) Pse5Ac7Ac at Ser-246 forms an H-bond with Lys-205 from domain D2 of the adjacent protofilament.

**SI Appendix, Fig. S7.**
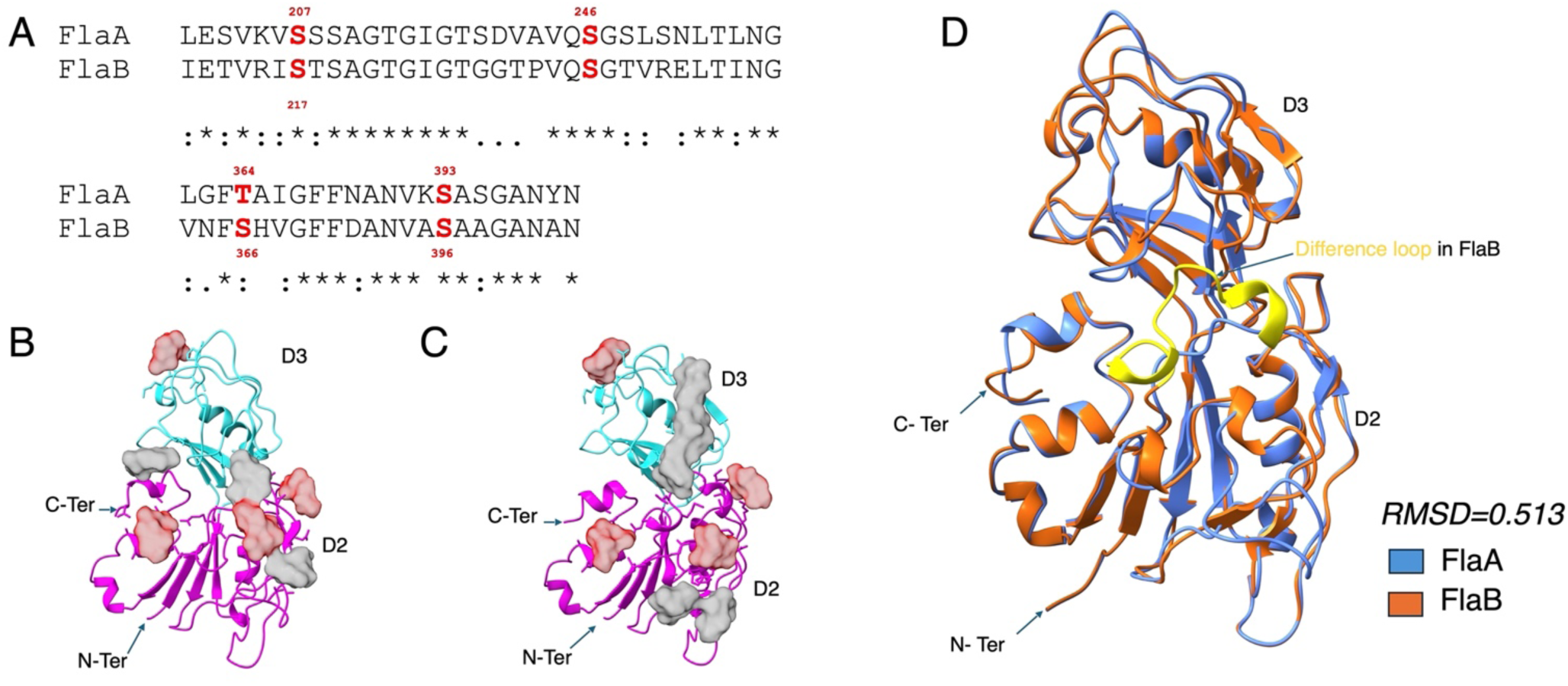
Comparison of conserved glycosylation sites in FlaA and FlaB. (**A**) Serine and threonine residues that were identified as being glycosylated in the *in-situ* structures of FlaA and FlaB are indicated in red. Amino acid residues surrounding the O-glycosylated residues that are identical in FlaA and FlaB are indicated with an asterisk (*) and residues with high similarity and lower similarity are indicated with a colon (:) and period (.), respectively. (**B**, **C**) Conserved glycosylation sites (red blob) highlighted in on FlaA and FlaB. Gray color blob represents non conserved glycosylation sites. (**D**) Structure alignment of domain D2 and D3 in FlaA and FlaB. FlaA and FlaB surface exposed domains are structurally similar with RMSD value of 0.513. The major difference between FlaA and FlaB lies in a loop that is present only in FlaB structure (colored yellow).

**SI Appendix, Fig. S8.**
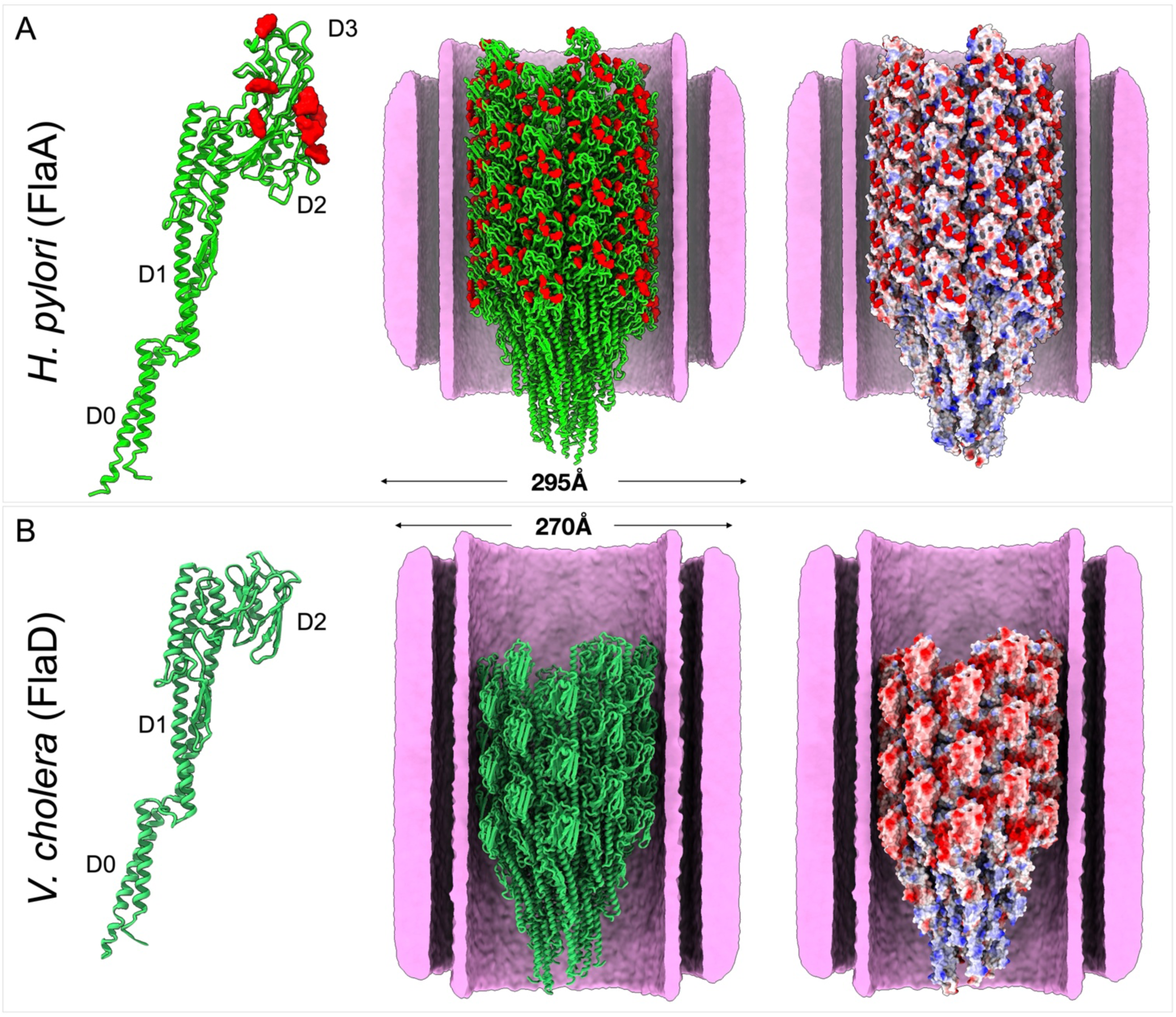
Comparison of sheathed flagella from *H. pylori* and *V. cholerae*. (**A**) Left panel: FlaA monomer model of *H. pylori*. Middle panel: FlaA filament model of *H. pylori*. Right panel: Surface charge property of the FlaA filament from *H. pylori.* Pseudaminic acids contribute to the negative charge on the FlaA filament surface in *H. pylori.* (**B**) Left panel: FlaD monomer model of *V. cholerae* (PDB ID: 9N8B). Middle panel: FlaD filament model of *V. cholerae*. Right panel: Surface charge property of the FlaD filament from *V. cholerae* is enriched by the acidic residues on D2 domain.

**SI Appendix, Table S1:** Cryo-EM data collection, analyses, and structure refinements.

| Parameters | FlaA<br>(middle region) | FlaA<br>(Tip region) | FlaB<br>(Near-hook region) |
| --- | --- | --- | --- |
| Magnification | 81000× |  |  |
| Voltage (kV) | 300 |  |  |
| Electron exposure (e-/Å<sup>2</sup>) | 70 |  |  |
| Defocus range (um) | -1.0 to -2.0 |  |  |
| Pixel size (Å) | 1.068 |  |  |
| Image processing |  |  |  |
| Helical rise (Å) | 65.403 | 65.403 | 65.400 |
| Helical twist (°) | 4.682 | 4.682 | 4.680 |
| Final number of particles | 228,410 | 275229 | 60029 |
| Map sharpening B factor (Å<sup>2</sup>) | -57.80 | -39.2 | -26.7 |
| Map resolution (Å) (FSC 0.143) | 2.65 Å | 3.19 Å | 3.22 Å |
| Refinement and model validation |  |  |  |
| Model composition | FlaA<br>(single flagellin) | FlaA<br>(single flagellin) | FlaB<br>(single flagellin) |
| Nonhydrogen atoms | 130383 | 84150 | 129228 |
| Number of amino acid residues | 16764 | 11176 | 16929 |
| Number of ligands | 231 | 154 | 297 |
| B factors (Å<sup>2</sup>)<br>Protein<br>Ligand | 33.01/208.03/76.04<br>72.34/148.88/103.67 | 37.62/215.28/88.26<br>57.27/167.72/100.42 | 54.81/271.39/112.03<br>103.24/217.61/157.34 |
| Root-mean-square deviations<br>Bond lengths (Å)<br>Bond angles (deg) | 0.005 (0)<br>0.480 (132) | 0.003 (0)<br>0.398 (35) | 0.002 (0)<br>0.450 (13) |
| MolProbity score | 2.14 | 2.74 | 2.45 |
| Clashscore | 20.14 | 32.64 | 39.28 |
| Rotamer outliers (%) | 1.53 | 3.47 | 1.86 |
| Ramachandran favored (%) | 96.79 | 95.28 | 96.99 |
| Ramachandran allowed (%) | 3.01 | 4.70 | 3.01 |
| Ramachandran outliers (%) | 0.20 | 0.02 | 0.01 |
| PDB ID | 9YGU | Not deposited | 9YH1 |
| EMDB ID | EMD-72941 | Not deposited | EMD-72948 |

## References

1. M. R. Amieva, E. M. El-Omar, Host-bacterial interactions in *Helicobacter pylori* infection. Gastroenterology 134, 306–323 (2008).

2. J. C. Atherton, M. J. Blaser, Coadaptation of *Helicobacter pylori* and humans: ancient history, modern implications. J Clin Invest 119, 2475–2487 (2009).

3. B. Linz et al., An African origin for the intimate association between humans and *Helicobacter pylori*. Nature 445, 915–918 (2007).

4. M. J. Blaser, *Helicobacter pylori*: microbiology of a ‘slow’bacterial infection. Trends Microbiol. 1, 255–260 (1993).

5. T. L. Cover, M. J. Blaser, *Helicobacter pylori* and gastroduodenal disease. Annu. Rev. Med. 43, 135–145 (1992).

6. E. J. Kuipers, *Helicobacter pylori* and the risk and management of associated diseases: gastritis, ulcer disease, atrophic gastritis and gastric cancer. Aliment Pharmacol Ther 11 Suppl 1, 71–88 (1997).

7. M. Bonis, C. Ecobichon, S. Guadagnini, M. C. Prevost, I. G. Boneca, A M23B family metallopeptidase of *Helicobacter pylori* required for cell shape, pole formation and virulence. Mol Microbiol 78, 809–819 (2010).

8. L. K. Sycuro et al., Peptidoglycan crosslinking relaxation promotes *Helicobacter pylori*’s helical shape and stomach colonization. Cell 141, 822–833 (2010).

9. D. R. Scott et al., The role of internal urease in acid resistance of *Helicobacter pylori*. Gastroenterology 114, 58–70 (1998).

10. K. A. Eaton, C. L. Brooks, D. R. Morgan, S. Krakowka, Essential role of urease in pathogenesis of gastritis induced by *Helicobacter pylori* in gnotobiotic piglets. Infect Immun 59, 2470–2475 (1991).

11. K. A. Eaton, D. R. Morgan, S. Krakowka, Motility as a factor in the colonisation of gnotobiotic piglets by *Helicobacter pylori*. J. Med. Microbiol. 37, 123–127 (1992).

12. K. M. Ottemann, A. C. Lowenthal, *Helicobacter pylori* uses motility for initial colonization and to attain robust infection. Infect Immun 70, 1984–1990 (2002).

13. J. Chu, J. Liu, T. R. Hoover, Phylogenetic distribution, ultrastructure, and function of bacterial flagellar sheaths. Biomolecules 10, 363 (2020).

14. T. Minamino, K. Imada, The bacterial flagellar motor and its structural diversity. Trends Microbiol 23, 267–274 (2015).

15. T. Fujii, T. Kato, K. Namba, Specific arrangement of alpha-helical coiled coils in the core domain of the bacterial flagellar hook for the universal joint function. Structure 17, 1485–1493 (2009).

16. F. A. Samatey et al., Structure of the bacterial flagellar hook and implication for the molecular universal joint mechanism. Nature 431, 1062–1068 (2004).

17. K. Rosinke, S. Tachiyama, J. Mrasek, J. Liu, T. R. Hoover, A *Helicobacter pylori* flagellar motor accessory is needed to maintain the barrier function of the outer membrane during flagellar rotation. PLoS Pathog 21, e1012860 (2025).

18. S. Tachiyama et al., FlgY, PflA, and PflB form a spoke-ring network in the high-torque flagellar motor of *Helicobacter pylori*. Proc Natl Acad Sci U S A 122, e2421632122 (2025).

19. X. Liu et al., Bacterial flagella hijack type IV pili proteins to control motility. Proc Natl Acad Sci U S A 121, e2317452121 (2024).

20. J. M. Botting et al., FlgV forms a flagellar motor ring that is required for optimal motility of *Helicobacter pylori*. PLoS One 18, e0287514 (2023).

21. E. Andersen-Nissen et al., Evasion of Toll-like receptor 5 by flagellated bacteria. Proc Natl Acad Sci U S A 102, 9247–9252 (2005).

22. A. T. Gewirtz et al., *Helicobacter pylori* flagellin evades toll-like receptor 5-mediated innate immunity. J Infect Dis 189, 1914–1920 (2004).

23. S. K. Lee et al., *Helicobacter pylori* flagellins have very low intrinsic activity to stimulate human gastric epithelial cells via TLR5. Microbes Infect 5, 1345–1356 (2003).

24. M. Kostrzynska, J. D. Betts, J. W. Austin, T. J. Trust, Identification, characterization, and spatial localization of two flagellin species in *Helicobacter pylori* flagella. J Bacteriol 173, 937–946 (1991).

25. C. Josenhans, A. Labigne, S. Suerbaum, Comparative ultrastructural and functional studies of Helicobacter pylori and Helicobacter mustelae flagellin mutants: both flagellin subunits, FlaA and FlaB, are necessary for full motility in *Helicobacter* species. J Bacteriol 177, 3010–3020 (1995).

26. M. Schirm et al., Structural, genetic and functional characterization of the flagellin glycosylation process in Helicobacter pylori. Mol Microbiol 48, 1579–1592 (2003).

27. P. Lertsethtakarn, K. M. Ottemann, D. R. Hendrixson, Motility and chemotaxis in *Campylobacter* and *Helicobacter*. Annu Rev Microbiol 65, 389–410 (2011).

28. A. Punjani, J. L. Rubinstein, D. J. Fleet, M. A. Brubaker, cryoSPARC: algorithms for rapid unsupervised cryo-EM structure determination. Nat Methods 14, 290–296 (2017).

29. E. J. O’Brien, P. M. Bennett, Structure of straight flagella from a mutant *Salmonella*. J Mol Biol 70, 133–152 (1972).

30. K. Yonekura, S. Maki-Yonekura, K. Namba, Complete atomic model of the bacterial flagellar filament by electron cryomicroscopy. Nature 424, 643–650 (2003).

31. J. Abramson et al., Accurate structure prediction of biomolecular interactions with AlphaFold 3. Nature 630, 493–500 (2024).

32. J. Jumper et al., Highly accurate protein structure prediction with AlphaFold. Nature 596, 583–589 (2021).

33. P. Emsley, K. Cowtan, Coot: model-building tools for molecular graphics. Acta Crystallogr D Biol Crystallogr 60, 2126–2132 (2004).

34. D. Liebschner et al., Macromolecular structure determination using X-rays, neutrons and electrons: recent developments in Phenix. Acta Crystallogr D Struct Biol 75, 861–877 (2019).

35. M. A. B. Kreutzberger, C. Ewing, F. Poly, F. Wang, E. H. Egelman, Atomic structure of the *Campylobacter jejuni* flagellar filament reveals how epsilon Proteobacteria escaped Toll-like receptor 5 surveillance. Proc Natl Acad Sci U S A 117, 16985–16991 (2020).

36. A. T. Franco et al., Delineation of a carcinogenic Helicobacter pylori proteome. Mol Cell Proteomics 8, 1947–1958 (2009).

37. H. Asakura et al., Helicobacter pylori HP0518 affects flagellin glycosylation to alter bacterial motility. Mol Microbiol 78, 1130–1144 (2010).

38. J. K. Chu et al., Loss of a cardiolipin synthase in *Helicobacter pylori* G27 blocks flagellum assembly. J Bacteriol 201, e00372–00319 (2019).

39. T. Minamino, K. Namba, Distinct roles of the FliI ATPase and proton motive force in bacterial flagellar protein export. Nature 451, 485–488 (2008).

40. H. S. Kim et al., The cell shape-determining Csd6 protein from *Helicobacter pylori* constitutes a new family of L,D-carboxypeptidase. J Biol Chem 290, 25103–25117 (2015).

41. L. K. Sycuro et al., Flow cytometry-based enrichment for cell shape mutants identifies multiple genes that influence *Helicobacter pylori* morphology. Mol Microbiol 90, 869–883 (2013).

42. J. A. Fuerst, Bacterial sheathed flagella and the rotary motor model for the mechanism of bacterial motility. J Theor Biol 84, 761–774 (1980).

43. W. Guo et al., Structures of the sheathed flagellum reveal mechanisms of assembly and rotation in *Vibrio cholerae*. Nature Microbiology 10, 3305–3314 (2025).

44. R. Einenkel et al., The structure of the complete extracellular bacterial flagellum reveals the mechanism of flagellin incorporation. Nat Microbiol 10, 1741–1757 (2025).

45. D. N. Mastronarde, SerialEM: a program for automated tilt series acquisition on Tecnai microscopes using prediction of specimen position. Microscopy and Microanalysis 9, 1182–1183 (2003).

46. I. A. Chen et al., The IMG/M data management and analysis system v.6.0: new tools and advanced capabilities. Nucleic Acids Res 49, D751–D763 (2021).

47. T. D. Goddard et al., UCSF ChimeraX: Meeting modern challenges in visualization and analysis. Protein Science 27, 14–25 (2018).

48. V. B. Chen et al., MolProbity: all-atom structure validation for macromolecular crystallography. Acta Crystallogr D Biol Crystallogr 66, 12–21 (2010).

49. N. P. Brown, C. Leroy, C. Sander, MView: a web-compatible database search or multiple alignment viewer. Bioinformatics 14, 380–381 (1998).

50. H. Ashkenazy et al., ConSurf 2016: an improved methodology to estimate and visualize evolutionary conservation in macromolecules. Nucleic Acids Res 44, W344–350 (2016).

